# Regulation of blood pressure by METTL3 via RUNX1b-eNOS pathway in endothelial cells

**DOI:** 10.1101/2023.12.11.571191

**Authors:** Yanhong Zhang, Xiaoxiao Yang, Mei Lan, Ze Yuan, Shuai Li, Yangping Liu, Cha Han, Ding Ai, Yang Yang, Yi Zhu, Bochuan Li

## Abstract

**Background:** Endothelial cells regulate vascular tone to control the blood pressure (BP) by producing both relaxing and contracting factors. Previously, we identified methyltransferase-like 3 (METTL3), a primary *N^6^*-methyladenosine (m^6^A) methyltransferase, as a key player in alleviating endothelial atherogenic progression. However, its involvement in BP regulation remains unclear.

**Methods:** To evaluate the role of METTL3 *in vivo*, mice with EC specific METTL3 deficiency (EC-*Mettl3^KO^*) with or without Ang II infusion were used to create a hypertensive model. Functional and MeRIP sequencing analysis were performed to explore the mechanism of METTL3-mediated hypertension.

**Results:** We observed a reduction in endothelial METTL3 activity by Ang II *in vitro* and *in vivo*. Endothelial METTL3-deficient mice exhibited higher BP than controls, both before and after Ang II infusion. Through m^6^A sequencing and functional analysis, we identified m^6^A modification of various RUNX1 monomers resulted in endothelial dysfunction. Mutations in the 3′UTR region of RUNX1b abolished its luciferase reporter activity, and enhanced eNOS promoter luciferase reporter activity with or without METTL3 overexpression. Overexpression of METTL3 by adeno-associated virus reduced Ang II-induced BP elevation.

**Conclusion:** This study reveals that METTL3 alleviates hypertension through m^6^A-dependent stabilization of *RUNX1b* mRNA, leading to upregulation of eNOS, thus underscoring the pivotal role of RNA transcriptomics in the regulation of hypertension.

## Introduction

*N*^6^-methyladenosine (m^6^A) is the most prevalent post-transcriptional modification of eukaryotic mRNAs^1^. This reversible modification is orchestrated by a multicomponent methyltransferase complex consisting of various methyltransferases including methyltransferase-like 3 (METTL3), METTL14, Wilms Tumor 1 Associated Protein (WTAP) and KIAA1429 (Virilizer). Conversely, demethylases such as fat mass and obesity-associated protein (FTO) and α-ketoglutarate-dependent dioxygenase alk B homolog 5 (ALKBH5) are responsible for its erasure^2^. As extensively documented, m^6^A on mRNAs plays a pivotal role in regulation of various cellular processes, including RNA stability, translation efficiency, RNA secondary structure, subcellular localization, alternative polyadenylation and splicing^3–5^. We previously reported that METTL3, the primary methyltransferase, alleviates endothelial atherogenic progression^6^. However, its role in hypertension remains unclear.

Hypertension is one of the most potent risk factors for cardiovascular morbidity and mortality and is characterized by compromised vascular function and a close connection with endothelial dysfunction^7^. Endothelial cells (ECs) form the inner lining of vascular beds and play a crucial role in the release of vasoactive factors including nitric oxide (NO). NO holds a pivotal position in the maintenance of vascular tone and normal BP^8^. In particular, NO serves as a crucial vasodilator, facilitating endothelium-dependent relaxation (EDR). Its production is intricately linked to the increased synthesis, modification and activity of endothelial NO synthase (eNOS)^9^. Achieving a delicate balance in the production of these vasoactive factors is paramount for the maintenance normal BP. However, the compensatory mechanisms underlying the regulation of this balance have yet to be fully elucidated.

RUNX1 (also known as AML1) is predominantly recognized for its role as a key transcriptional core-binding factor, influencing cell proliferation, differentiation, and survival during hematopoietic development^10^. Its interaction with the partner core binding factor beta (CBFβ) is essential for regulating the transcription of multiple genes^11^. Notably, RUNX1 has been identified as an emerging therapeutic target in cardiovascular disease^12^. Studies on RUNX1 null models have revealed a complete absence of definitive hematopoiesis, abnormal vasculature development, and reduced endothelial sprouting^13,14^. Intriguingly, in the context of cardiovascular pathology, RUNX1 exacerbates pathological cardiac hypertrophy by activating p53 in response to Ang II^15^. This dual role underscores the complexity of RUNX1 function in different cellular contexts and warrants further exploration.

In this study, we employed tamoxifen-inducible endothelial-specific *Mettl3*-deficient (EC-*Mettl3^KO^*) mice, as previously described^6^, to investigate the regulatory role of Mettl3-mediated m^6^A in hypertension. Our data revealed that the downregulation of METTL3 and subsequent hypomethylation contributed to Ang II-induced endothelial dysfunction and hypertension, both *in vivo* and *in vitro.* Notably, we identified RUNX1b as a pivotal regulatory target in METTL3-dependent EC activation. These findings shed light on the critical mechanism involving m^6^A modifications in BP regulation.

### Methods

All supporting data are available within the article and its Supplemental Material. Detailed descriptions of experimental methods of the current study are provided in the Supplemental Material.

### Data Availability

RNA-seq and MeRIP-seq data generated in this study have been deposited to the Genome Sequence Archive in BIG Data Center under accession number GSE248592.

## Results

### METTL3 is decreased in an Ang II-induced hypertensive model

To elucidate the role of m^6^A modification in Ang II-induced hypertension, we first established a hypertensive mouse model using a 3-day infusion of Ang II, which resulted in elevated systolic BP (SBP) and diastolic BP (DBP) (Figure S1A-B). To investigate the impact of Ang II on m^6^A modulators, we assessed the expression of key methyltransferase components, METTL3, METTL14, WTAP, and Virilizer, in the intima and media of the aorta. Western blot analysis revealed higher expression of METTL3 in the intima than in the media, which significantly decreased after Ang II stimulation, with no observable changes in other m^6^A modulators (Figure 1A-B and Figure S1C-D). Concurrently, phosphorylated and total eNOS (p-eNOS and t-eNOS) in the intima, along with serum NO levels, decreased after Ang II stimulation (Figure 1A, C-D). Interestingly, METTL3 levels were more abundant in the mesenteric arteries (MA) compared to the aorta and left common carotid arteries (LCA), and were significantly reduced by Ang II (Figure S1E-F). Furthermore, Ang II exhibited time- and dose-dependent downregulation of both METTL3 and p-eNOS protein levels, without affecting other m^6^A modulators (Figure 1E-H). Collectively, METTL3 expression decreased in the Ang II-induced hypertensive model.

**Figure 1.**
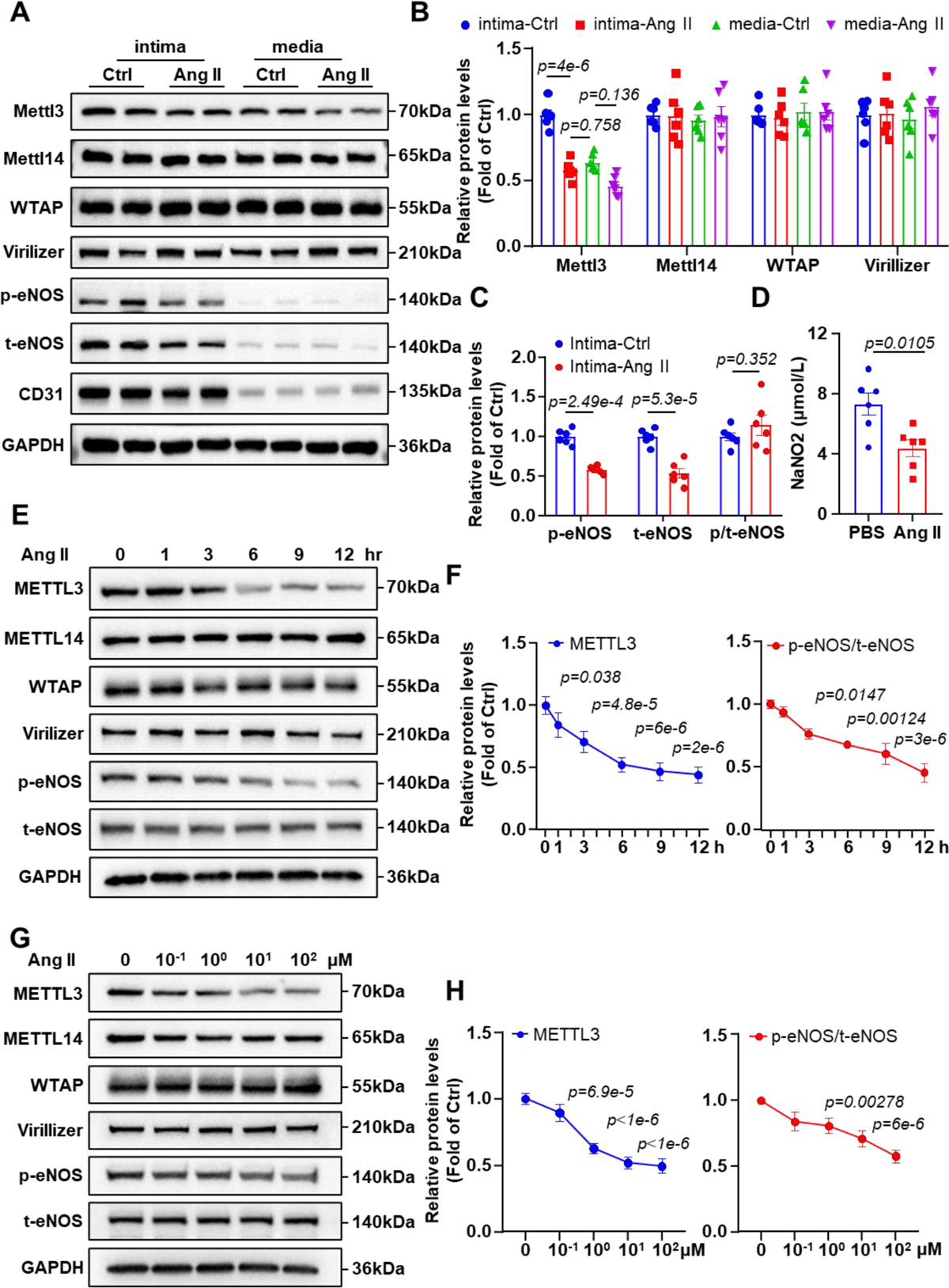
METTL3 is decreased in an Ang II-induced hypertensive model. **(A-B)** Protein was extracted from the aortic intima and media of male C57BL/6 mice induced by Ang II (490 ng/kg/min) for 3 days. Mice with normal blood pressure as a control. **(A)** Western blot analysis of Mettl3, Mettl14, WTAP, Virilizer, p-eNOS, t-eNOS, CD31 and GAPDH. (**B-C**) Quantification of the protein expression in (A). The data are presented as means ± the SEM, *p < 0.05, ns, not significant (two-way ANOVA with Bonferroni multiple comparison post hoc test). n=6 mice. **(D)** The content of nitric oxide (NO) in the plasma from Ctrl mice and Ang II induced mice, the data are shown as means ± the SEM, *p < 0.05 (Student’s *t* test). n=6 mice. **(E)** Western blot analysis of METTL3, METTL14, WTAP, Virilizer, p-eNOS, t-eNOS and GAPDH in human umbilical vein endothelial cells (HUVECs) treated with Ang II (10 nmol/L) for 0, 1, 3, 6, 9 and 12 h. **(F)** Quantification of the protein expression in (E). The data are presented as means ± the SEM, *p < 0.05 (one-way ANOVA with Bonferroni multiple comparison post hoc test). n=6. **(G)** Western blot analysis of METTL3, METTL14, WTAP, Virilizer, p-eNOS, t-eNOS and GAPDH in HUVECs treated with 0, 0.1, 1, 10, 100 nmol/L Ang II for 6 h. **(H)** Quantification of the protein expression in (G). The data are shown as means ± the SEM, *p < 0.05 (one-way ANOVA with Bonferroni multiple comparison post hoc test). n=6.

### Endothelial specific METTL3 deficiency increases BP via inhibition of eNOS

Next, to elucidate the impact of METTL3 deficiency on endothelial dysfunction and BP regulation, we utilized tamoxifen-inducible endothelial-specific *Mettl3*-deficient (EC*-Mettl3^KO^*) mice, as previously described ^6^. EC*-Mettl3^KO^*mice exhibited increased SBP and DBP at baseline and in response to Ang II compared to control mice (Figure 2A-B). The vasoconstrictive function remained unchanged between the 2 groups (Figure S2A-B). Disruption of METTL3 in ECs markedly impaired acetylcholine (ACh)-induced EDR in second-order MAs when compared with control mice (Figure 2C). This impairment was specific to endothelial dysfunction, as it did not affect endothelium-independent relaxation with sodium nitroprusside or vasoconstrictive responses to phenylephrine (Figure S2C-D). Further examination of EDR by treating the arteries with the nitric oxide synthase inhibitor L-NG-nitroarginine methyl ester (L-NAME), revealed reduced ACh-dependent relaxation, abolishing the difference in EDR between the 2 genotypes (Figure 2C). Moreover, METTL3 and total eNOS protein levels were higher in the mesenteric arteries than in the aorta and were significantly reduced by the disruption of METTL3 in ECs (Figure 2D-F and Figure S2E-F). Next, we evaluated NO generation in the aorta and MA using the electron paramagnetic resonance (EPR) spin-trapping reagent Fe(DETC)_2_, which indicated a decrease in the characteristic NO-Fe(DETC)_2_ EPR triplet in EC*-Mettl3^KO^* mice compared to that in control mice (Figure 2G-H). These results were consistent with the almost complete disappearance of METTL3 and t-eNOS expression upon METTL3 knockout in ECs (Figure S2G-H). These results suggest that NO-derived vasodilation plays a critical role in endothelial function induced by endothelial METTL3 deficiency in BP regulation.

**Figure 2.**
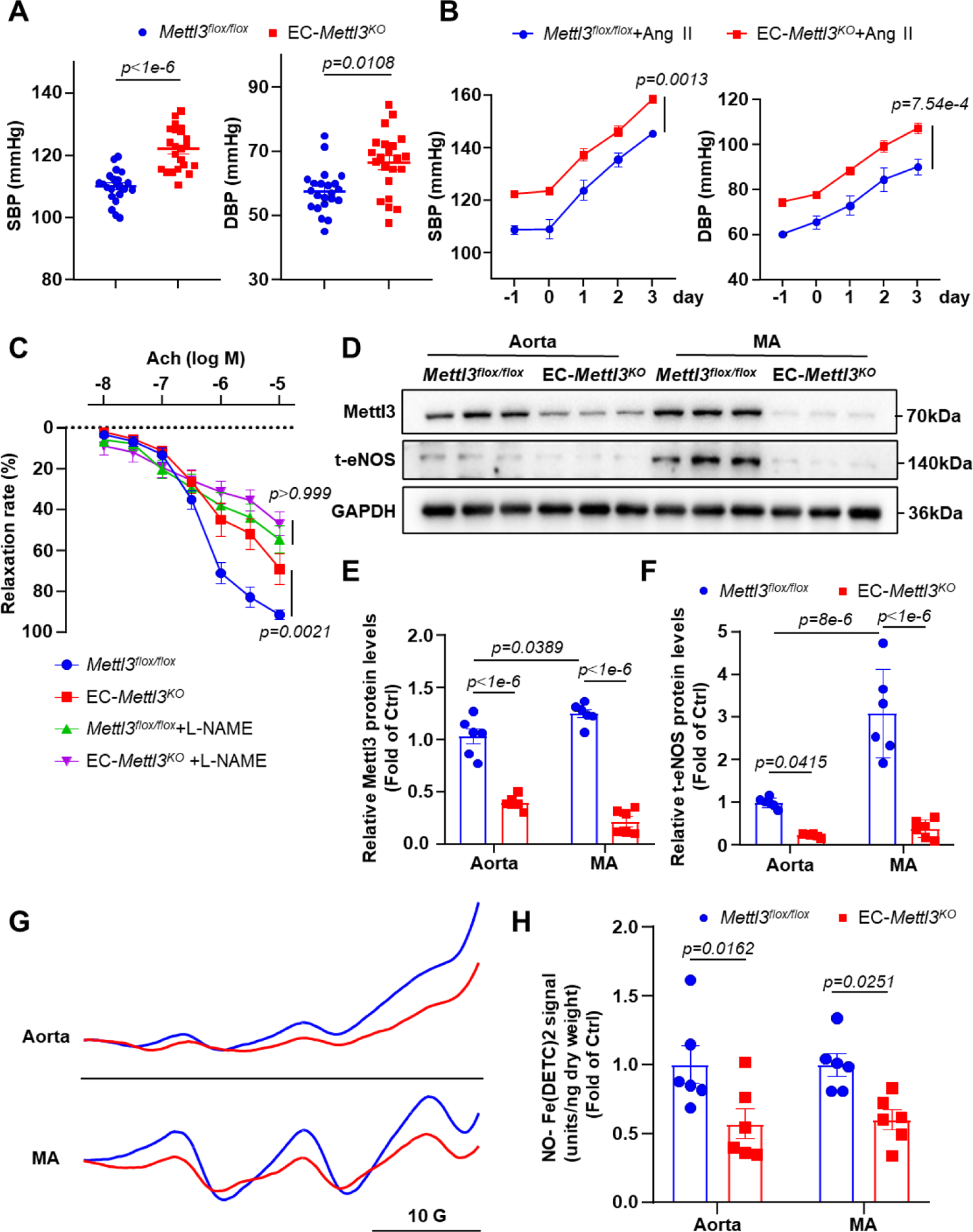
Endothelial specific METTL3 deficiency increases BP via inhibition of eNOS. **(A)** Noninvasive tail-cuff monitoring of SBP and DBP in 8-week-old *Mettl3^flox/flox^* and EC*-Mettl3^KO^* mice. The data are presented as means ± the SEM, *p < 0.05 (Student’s *t* test). n=22 mice. **(B)** SBP and DBP of *Mettl3^flox/flox^* and EC*-Mettl3^KO^* mice were infused with Ang II (490 ng/kg/min) for 3 days. The data are shown as means ± the SEM, *p < 0.05 (two-way ANOVA with Bonferroni multiple comparison post-test). n=6 mice. **(C)** MAs from *Mettl3^flox/flox^*and EC*-Mettl3^KO^* mice were pretreated with or without L-NAME for 30 mins, EDR in response to ACh was assessed. The data are presented as means ± the SEM, *p < 0.05, ns, not significant (two-way ANOVA with Bonferroni multiple comparison post-test). n=8 mice. **(D-F)** Protein was extracted from the Aorta and MA of 8-week-old *Mettl3^flox/flox^*and EC*-Mettl3^KO^* mice. **(D)** Western blot analysis of the expression of METTL3, t-eNOS and GAPDH in tissue lysates of Aorta and MA. **(E-F)** Quantification of the protein expression in (D). The data are shown as means ± the SEM, *p < 0.05 (two-way ANOVA with Bonferroni multiple comparison post hoc test). n=6 mice. **(G-H)** Representative EPR spectra and quantification of NO-Fe (DETC)_2_ signals from Aorta and Mesentery arteries of *Mettl3^flox/flox^*and EC*-Mettl3^KO^* mice. The data are presented as means ± the SEM, *p < 0.05 (Student’s *t* test). n=6 mice.

### METTL3 mediates methylation of RUNX1

As a core subunit of the m^6^A methyltransferase complex, the observed downregulation of METTL3 expression in response to Ang II suggests the potential regulation of m^6^A modification. We first conducted m^6^A-specific methylated RNA immunoprecipitation combined with high-throughput sequencing (MeRIP–seq) to compare the m^6^A landscape between Ang II-treated samples with or without METTL3 overexpression. We identified 9067 and 20119 m^6^A peaks in Ang II and Ang II with METTL3 overexpression conditions, respectively. The identified m^6^A motifs were significantly enriched in CGACG motif (Figure 3B) and were abundant in coding regions (CDSs), 3′ untranslated regions (3′UTRs), and near stop codons (Figure 3A and Figure S3A). Further, we screened thousands of dysregulated m^6^A peaks in the 2 groups (Figure S3B), and the downregulated genes in input RNA-seq were enriched in TNFα and cytokine-cytokine receptor interaction pathways (Figure S3C, right panel). Taken together, we found that METTL3 appears to mediate dynamic changes in the m^6^A landscape between Ang II and Ang II in association with METTL3 overexpression.

**Figure 3.**
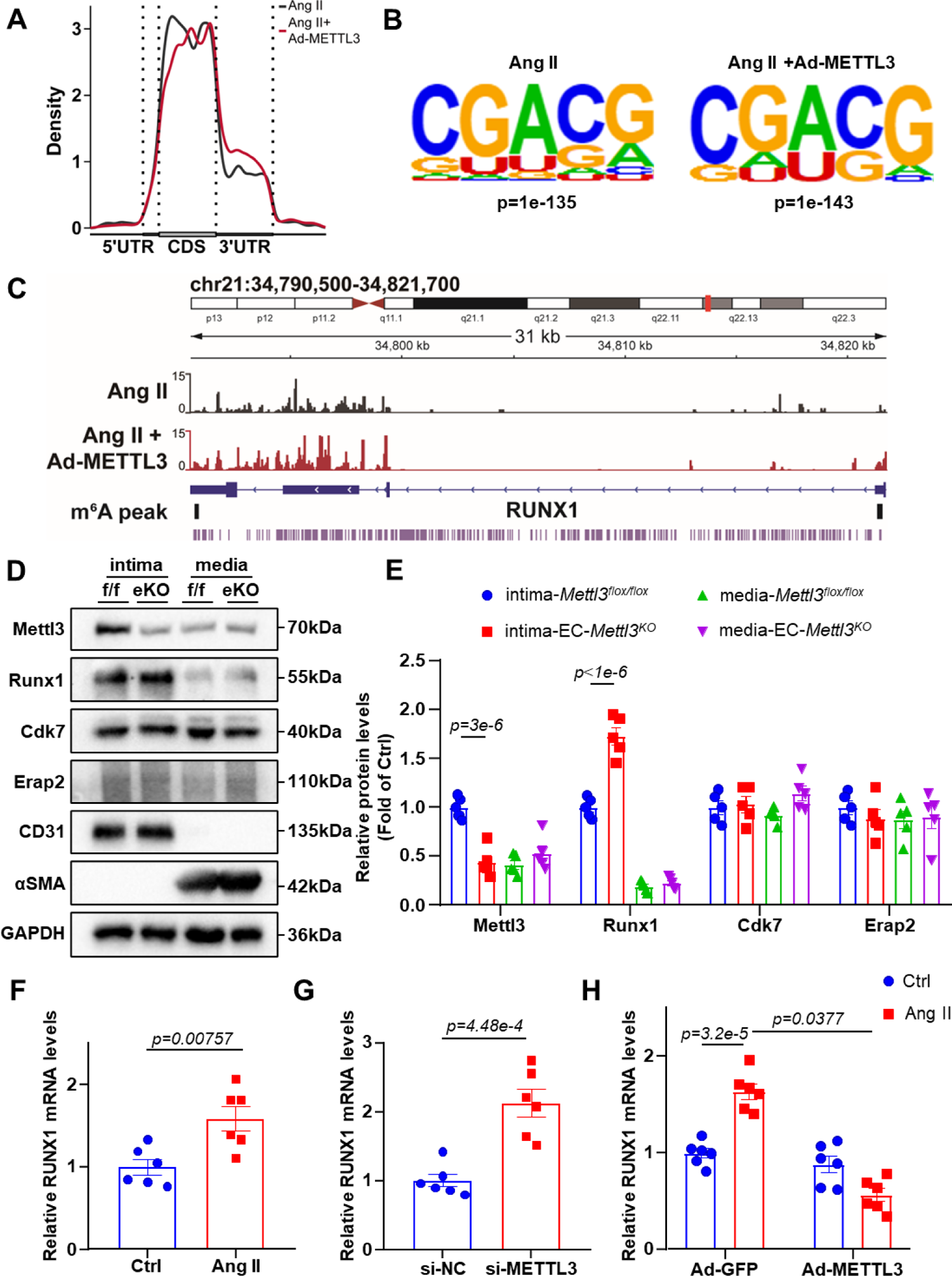
METTL3 mediates methylation of RUNX1 HUVECs were treated by Ang II (10 nmol/L) with or without adenovirus overexpression of METTL3 for MeRIP-seq analysis. **(A)** Distribution of m^6^A peaks along the 5’UTR, CDS, and 3’UTR regions of mRNA. (**B)** m^6^A motif identified from HUVECs of mRNA. **(C)** Integrative genomics viewer tracks displaying the results of m^6^A-IP vs. Input read distributions in RUNX1. m^6^A peak, high confidence region. **(D-E)** Protein was extracted from the aortic intima and media of *Mettl3^flox/flox^* (f/f) and EC*-Mettl3^KO^* (eKO) mice. **(D)** Western blot analysis of Mettl3, Runx1, Cdk7, Erap2, CD31, αSMA and GAPDH in tissue lysates of aortic intima and media. **(E)** Quantification of the protein expression in (D). The data are presented as means ± the SEM, *p < 0.05 (two-way ANOVA with Bonferroni multiple comparison post hoc test). n=5 mice. **(F-G)** HUVECs were treated by Ang II (10 nmol/L) for 6 hours or infected with si-METTL3 for 24 hours. Quantification of the mRNA expression of *RUNX1*. The data are shown as means ± the SEM, *p < 0.05 (Student’s *t* test). n=6. **(H)** HUVECs treated by Ang II (10 nmol/L) with or without adenovirus overexpression of METTL3. Quantification of the mRNA expression of *RUNX1*. The data are presented as means ± the SEM, *p < 0. 05 (two-way ANOVA with Bonferroni multiple comparison post hoc test). n=6.

As reported previously, m^6^A modification plays an important role in regulating RNA abundance. To investigate the molecular mechanisms of m^6^A in endothelial activation, we analyzed the regulators of methylation-upregulated but expression-downregulated genes. While exploring the methylation status of *NOS3*, the gene encoding eNOS, which is a core downstream target of METTL3, we found no direct regulation of METTL3-m^6^A methylation (Figure S3D). The m^6^A prediction also confirmed the low confidence of *NOS3* in SRAMP (a sequence-based *N^6^*-methyladenosine modification site predictor). As eNOS is transcriptionally regulated by METTL3, we identified RUNX1^16^, CDK7^17^, and ERAP2^18^ as 3 potential key regulators participating in Ang II-induced hypertension, which was further confirmed by the increased IP/input ratio (Figure 3C and Figure S3E-F). Among these, only Runx1 was specifically expressed in the vascular intima, with increased expression in EC-*Mettl3^KO^*mice compared to control mice. As controls, Cdk7 and Erap2 were widely expressed in the vascular tissues without significant difference (Figure 3D-E). The GEO database corroborated the increased enrichment of Runx1 in response to Ang II treatment (Figure S3G-I). We also observed a significant increase in *RUNX1* mRNA levels in response to Ang II and deletion of METTL3. Overexpression of METTL3 markedly rescued the increased *RUNX1* mRNA levels induced by Ang II (Figure 3F-H). Similar trends were observed *in vivo* and *in vitro*, confirming that Ang II-induced EC activation was mediated by m^6^A modification on RUNX1 (Figure S4A-F).

### RUNX1/eNOS axis signaling is involved in METTL3 deficiency-induced EC dysfunction

In order to validate whether eNOS is a direct target gene of RUNX1, we conducted a comprehensive promoter sequence analysis, identifying 2 RUNX1 binding sites in the promoter region of the eNOS gene (Figure 4A and Table S1). Subsequently, chromatin immunoprecipitation (ChIP) assays confirmed the direct connection between RUNX1 and eNOS promoter (Figure 4B-C). To further elucidate the functional relationship between RUNX1 and eNOS, we measured EDR by treating mesenteric arteries with L-NAME and the selective RUNX1-CBFβ interaction inhibitor, Ro5-3335. METTL3 deficiency-induced dysfunction of ACh-dependent vasodilation was successfully reversed by preincubation with Ro5-3335, while L-NAME abolished the difference in EDR between the 2 genotypes in the presence of Ro5-3335 (Figure 4D). These findings strongly suggest that METTL3 mediates eNOS-dependent EDR through the regulation of RUNX1. Western blot analysis further revealed that Ro5-3335 significantly downregulated RUNX1 levels but concurrently restored the expression of eNOS levels under METTL3 knockdown in ECs (Figure 4E-F). METTL3 was used as a marker of knockout efficiency (Figure 4G). Ro5-3335 markedly upregulated NO levels when METTL3 was knocked down in ECs (Figure 4H). These results provide compelling evidence that eNOS is a novel direct downstream target gene of RUNX1 and that inhibition of RUNX1 ameliorates the decreased eNOS expression caused by *METTL3* knockout.

**Figure 4.**
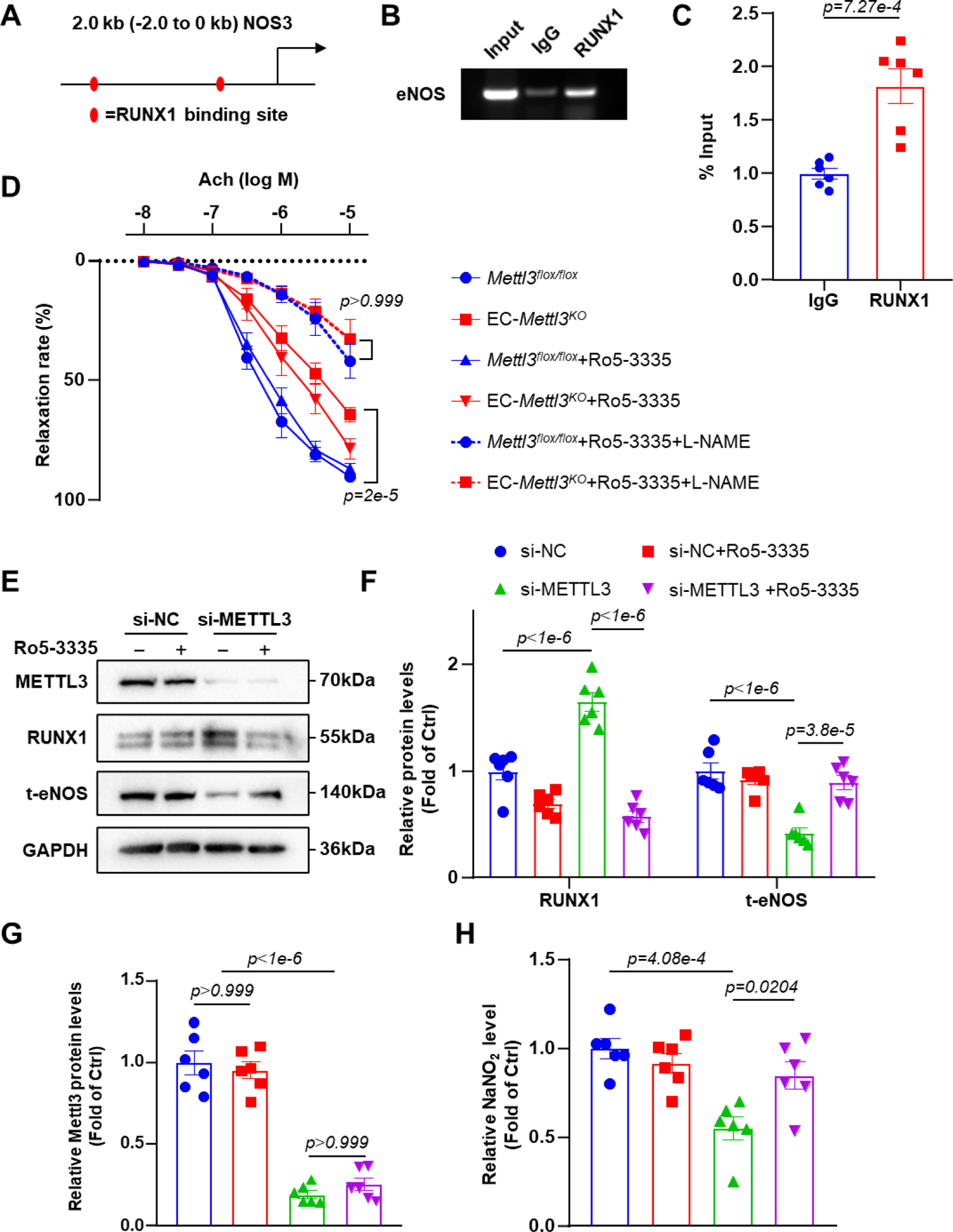
RUNX1/eNOS axis signaling is involved in METTL3 deficiency-induced EC dysfunction. **(A)** Sequence analysis of RUNX1 binding sites (red oval) in the promoter region of NOS3 gene. **(B)** Chromatin immunoprecipitation (ChIP) assay of RUNX1 and the promoter of NOS3 gene with gel images. **(C)** Chromatin immunoprecipitation (ChIP) assay of RUNX1 and the promoter of NOS3 gene with qPCR analysis. The data are presented as means ± the SEM, *p < 0.05 (Student’s *t* test). n=6. **(D)** MAs isolated from *Mettl3^flox/flox^* and EC*-Mettl3^KO^* mice were pretreated with Ro5-3335 (10 umol/L) or L-Name (100 umol/L) in response to ACh. The data are shown as means ± the SEM, *p < 0.05 (two-way ANOVA with Bonferroni multiple comparison post hoc test). n=6 mice. **(E-H)** HUVECs were infected with siRNA of METTL3 for 24 hours and treated with or without Ro5-3335 (10 μmol/L) during the last 6 hours. **(E)** Western blot analysis of METTL3, RUNX1, t-eNOS and GAPDH. **(F-G)** Quantification of the protein expression in (E). The data are presented as means ± the SEM, *p < 0.05, ns, not significant. (two-way ANOVA with Bonferroni multiple comparison post-test). n=6. **(H)** The content of nitric oxide (NO) in the HUVECs lysis, the data are shown as means ± the SEM, *p < 0.05 (two-way ANOVA with Bonferroni multiple comparison post-test). n=6.

### Exogenous METTL3 supplementation mitigates Ang II-induced hypertension

To elucidate the function of METTL3 in Ang II-induced hypertension, we examined the BP levels in adeno-associated virus-overexpressing METTL3 (AAV9-METTL3 OE)-infected mice with or without Ang II pumping for 3 days. Notably, METTL3 overexpression attenuated Ang II-induced BP elevation (Figure 5A). As expected, AAV9-METTL3 OE reduced the increase in RUNX1 protein levels by Ang II, as shown by immunofluorescence staining and western blot analysis (Figure 5B-E). Concurrently, AAV9-METTL3 OE reversed the reduction in eNOS protein levels induced by Ang II (Figure S5A-B). The efficiency of METTL3 overexpression was confirmed using immunofluorescence staining (Figure S5C-D). In addition, EPR detection of NO confirmed that AAV9-METTL3 OE rescued the decrease in NO release in EC-*Mettl3^KO^* mice (Figure 5F-G). These data indicated that endothelial METTL3 is a pivotal regulator of EDR and is dependent on the RUNX1/eNOS pathway *in vivo*.

**Figure 5.**
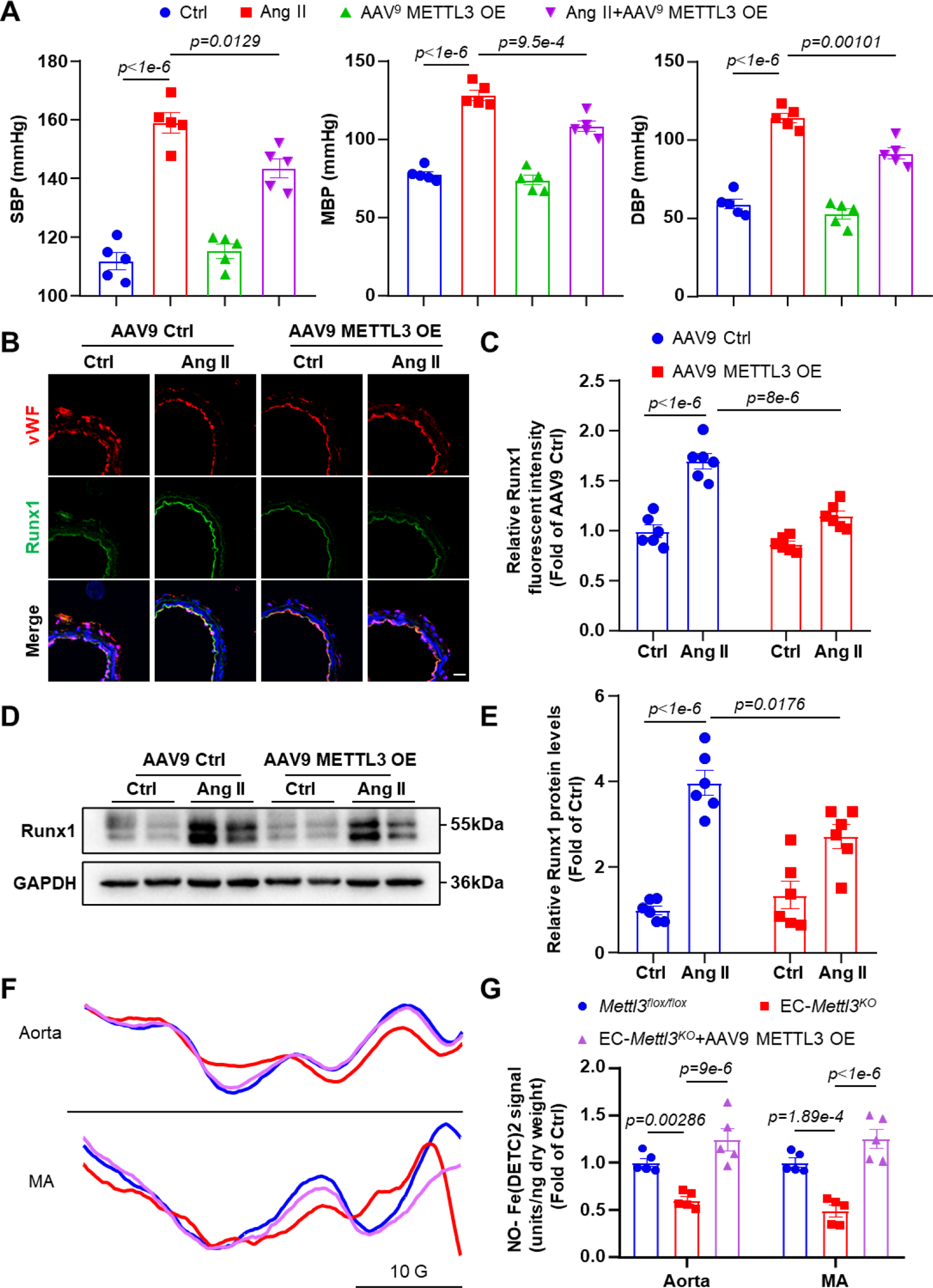
Exogenous METTL3 supplementation mitigates Ang II-induced hypertension. **(A)** Noninvasive tail-cuff monitoring of SBP, MBP and DBP in male C57BL/6 mice induced by Ang II (490 ng/kg/min) for 3 days with or without Adeno-associated virus overexpression of METTL3 (AAV9^endo^ METTL3). The data are presented as means ± the SEM, *p < 0.05 (two-way ANOVA with Bonferroni multiple comparison post hoc test). n=5 mice. **(B-C)** Male C57BL/6 mice induced by Ang II (490ng/kg/min) for 3 days with or without Adeno-associated virus overexpression of METTL3. **(B)** MA underwent immunofluorescence staining for indicated proteins. Scale bar, 20μm. **(C)** Quantification of the relative fluorescence intensity in (B). The data are shown as means ± the SEM, *p < 0.05 (two-way ANOVA with Bonferroni multiple comparison post-test). n=6 mice. **(D)** Protein was extracted from the MA. Western blot analysis of Runx1 and GAPDH. **(E)** Quantification of the protein expression in (D). The data are presented as means ± the SEM, *p < 0.05 (two-way ANOVA with Bonferroni multiple comparison post hoc test). n=6 mice. **(F-G)** Representative EPR spectra and quantification of NO-Fe (DETC)_2_ signals from Aorta and MA of *Mettl3^flox/flox^*, EC*-Mettl3^KO^* and EC*-Mettl3^KO^* with Adeno-associated virus overexpression of METTL3 mice. The data are shown as means ± the SEM, *p < 0.05 (one-way ANOVA with Bonferroni multiple comparison post hoc test). n=5 mice.

### RUNX1b is mainly responsible for METTL3-induced eNOS downregulation and endothelial dysfunction

RUNX1 exhibits a complex expression pattern, regulated at the levels of transcription, splicing and translation^19^. RUNX1 has four variants: RUNX1a, RUNX1b, RUNX1c, and RUNX1△N, and the various RUNX1 isoforms are functionally different^20^. In-depth analysis revealed that RUNX1b was prominently influenced by knockout of METTL3 in ECs; RUNX1a exhibited a lower usage rate than RUNX1b when treated with Ang II and was partially rescued by METTL3 overexpression in RNA-seq (Figure 6A, left and middle panels). Based on MeRIP-seq, RUNX1a exhibited a slightly higher usage rate than RUNX1b (Figure 6A, right panel). RUNX1c expression was almost undetectable across all experimental conditions. Subsequently, both RUNX1a and RUNX1b became hypermethylated following METTL3 overexpression (Figure 3C and 6B). To investigate the regulatory role of METTL3-mediated m^6^A in the decay of *RUNX1a/b* mRNA, we measured *RUNX1a/b* mRNA levels in ECs after treatment with the transcriptional inhibitor actinomycin D. Compared to the GFP control, METTL3 overexpression significantly decreased the remaining *RUNX1b* mRNA levels, owing to accelerated mRNA decay in the presence of functional m^6^A modification (Figure 6C). Conversely, *RUNX1b* mRNA exhibited a slower decay rate in response to METTL3 knockdown and Ang II treatment (Figure 6D-E). Interestingly, RUNX1a was not degraded by METTL3 (Figure S6A-B). Further investigation into the mechanism of RUNX1 regulation by METTL3 revealed that RUNX1a was alternatively spliced at exon 7, which is the best-known splicing site of RUNX1a^21^, based on RNA-seq and meRIP-seq (data not shown). As we found that the 3′UTR of RUNX1b and CDSs of RUNX1a are key regions for regulating m^6^A modification, we conducted mutational analysis on the potential m^6^A motifs. Specifically, we mutated four nearby potential RUNX1a m^6^A motifs from A to T, and three RUNX1b mutants: proximal mutants (RUNX1b mut2), distal mutants (RUNX1b mut3) and a combination of all RUNX1 potential mutants (RUNX1b mut1). Despite the successful abolishment of increased RUNX1a in luciferase activity, they did not affect the increase in eNOS promoter luciferase activity caused by overexpression of METTL3 (Figure S6C-D). Interestingly, mutations of the proximal (mut2) and distal (mut3) sites of RUNX1b exhibited entirely opposite effects on the luciferase activity (Figure 6F). Specifically, RUNX1b mut2 enhanced eNOS luciferase activity with or without METTL3 overexpression, whereas RUNX1b mut3 did not have a discernible effect (Figure 6G). Furthermore, knockdown of RUNX1b significantly rescued the down-regulation of eNOS and NO production by disruption of METTL3. METTL3 protein and RUNX1b mRNA levels were used as markers of knockdown efficiency (Figure 6H-I and Figure S6E-F). Collectively, these findings strongly suggest that RUNX1b is the key downstream of m^6^A modification by METTL3 in the regulation of eNOS promoter activity and endothelial dysfunction.

**Figure 6.**
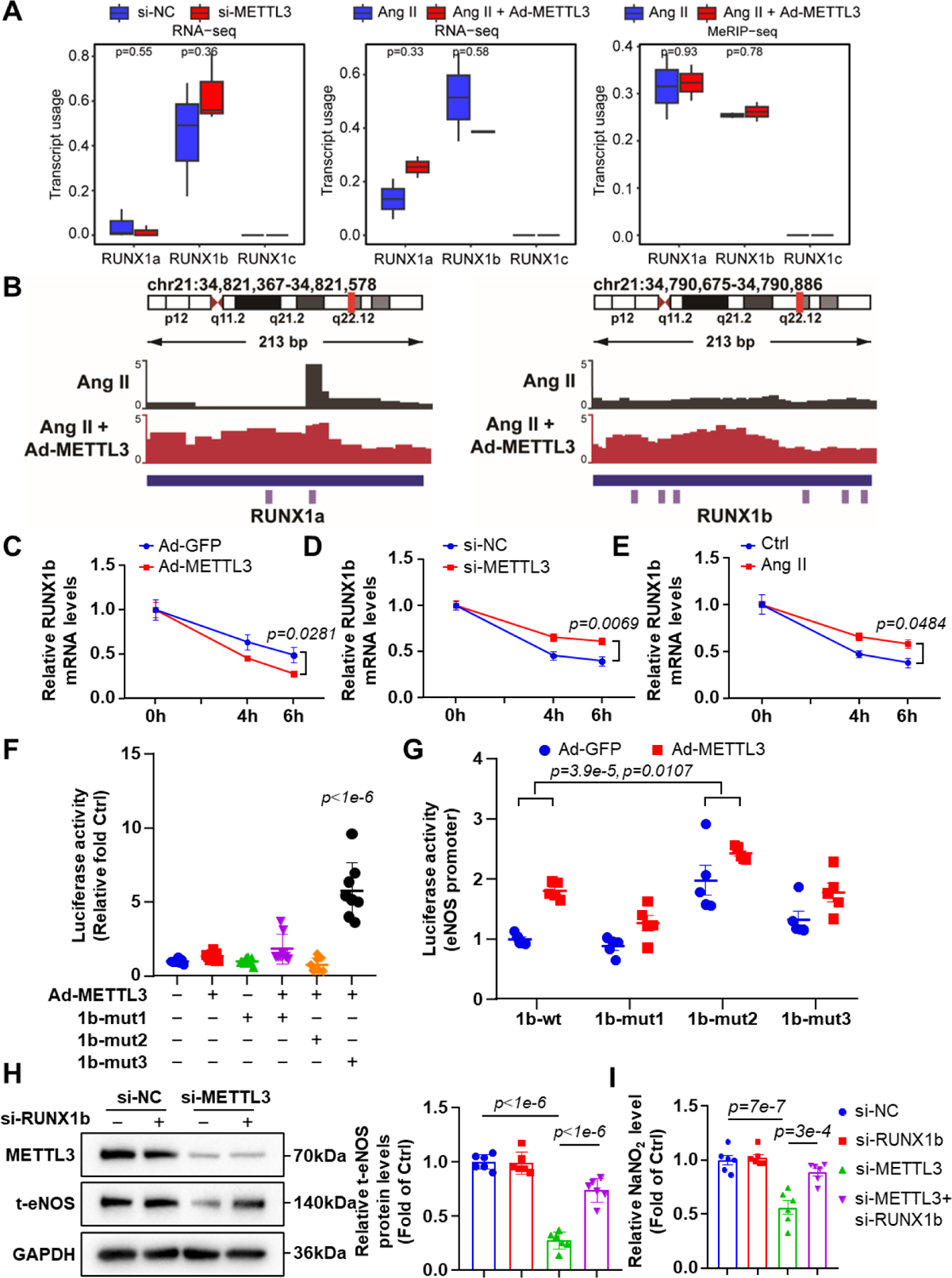
RUNX1b is mainly responsible for METTL3-induced eNOS downregulation and endothelial dysfunction. **(A)** HUVECs were treated by knockdown of METTL3 or Ang II (10 nmol/L) with or without adenovirus overexpression of METTL3. Usage of RUNX1a/b/c in RNA-seq and MeRIP-seq. **(B)** Integrative genomics viewer tracks displaying the results of m^6^A-IP vs. Input read distributions in *RUNX1a* and *RUNX1b* mRNA. **(C-E)** HUVECs were treated with actinomycin D for 4 and 6 hr. qPCR analysis showing RUNX1b mRNA degradation upon METTL3-overexpression (C); si-METTL3 (D); and Ang II (10 nmol/L) treatment (E). The data are presented as means ± the SEM, *p < 0.05 (Student’s *t* test). n=5 (C), n= 6 (D-E). **(F-G)** Relative activity of eNOS promoter luciferase in the wild-type or mutant RUNX1a or RUNX1b transfected in K293 cells treated with METTL3-overexpressing adenovirus. The data are shown as means ± the SEM, *p < 0.05, ns, not significant (two-way ANOVA with Bonferroni multiple comparison post-test). n=8 (F), n=5 (G). **(H-I)** HUVECs were infected with knockdown of METTL3 with or without knockdown of RUNX1b for 24 hours. Western blot analysis of METTL3, RUNX1, t-eNOS and GAPDH. The data are presented as means ± the SEM, *p < 0.05 (two-way ANOVA with Bonferroni multiple comparison post-test). n=6. **(I)** The content of nitric oxide (NO) in the HUVECs lysis, the data are shown as means ± the SEM, *p < 0.05 (two-way ANOVA with Bonferroni multiple comparison post-test). n=6.

## Discussion

Our findings underscore the pivotal role of RNA m^6^A modification in the regulation of endothelial activation and hypertensive response to Ang II. First, the reduction in m^6^A modification levels, concomitant with the downregulation of METTL3 in response to Ang II, sets the stage for a cascade of events. Secondly, endothelial-specific METTL3 deficiency induced a significant reduction in eNOS, a central modulator of hypertension. Further, RUNX1 negatively regulates the transcription of eNOS by binding to its promotor, and RUNX1 inhibition prevents the development of hypertension. Finally, sequencing and functional analyses unveil that m^6^A modification of the *RUNX1b* 3′UTR contributes to endothelial dysfunction, suggesting a novel therapeutic avenue for hypertension. Together, our findings demonstrate that METTL3 and m^6^A modifications could alleviate endothelial activation and hypertension through accelerated degradation of *RUNX1b* mRNA in response to Ang II (Figure 7).

**Figure 7.**
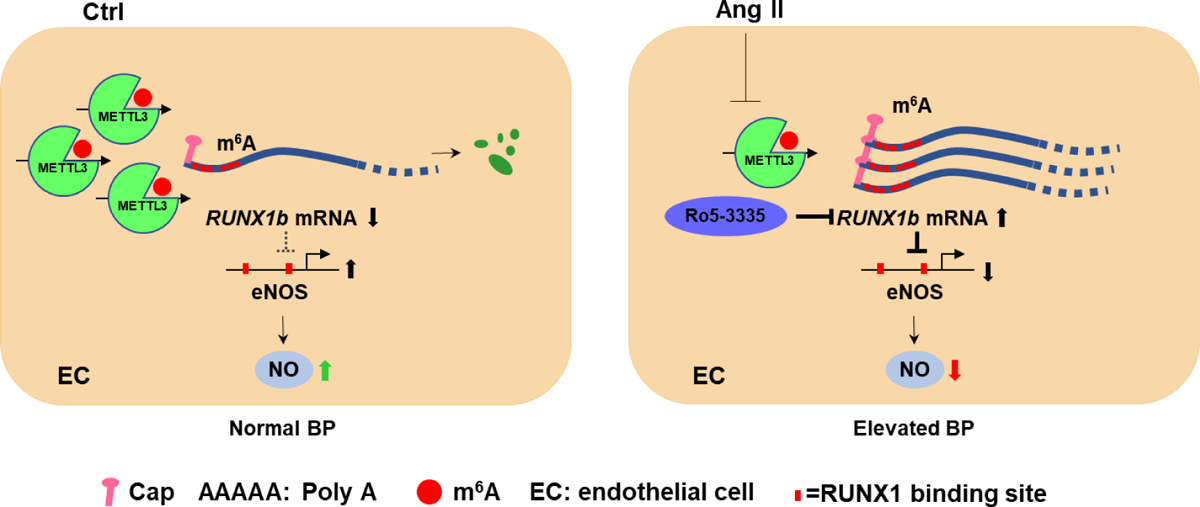
A model illustrating the protective role of METTL3 and m^6^A in alleviating hypertension. At static ECs, METTL3 catalyzes the formation of m^6^A modification, thus facilitates *RUNX1b* degradation to maintain EC homeostasis. In response to Ang II, decreased METTL3 fails to modulate *RUNX1b* degradation, leading to RUNX1 signaling activation and elevating blood pressure.

Growing evidence supports the involvement of aberrant m^6^A methylation in various cardiovascular diseases, including cardiac hypertrophy, heart failure, vascular calcification, atherosclerosis, and pulmonary hypertension^22,23^. Dorn et al. demonstrated that enhanced m^6^A RNA methylation resulted in compensated cardiac hypertrophy, while diminished m^6^A correlated with signs of eccentric cardiomyocyte remodeling and failure, highlighting the importance of m^6^A in the maintenance of cardiac homeostasis^24^. Other studies have implicated m^6^A in the stability of circRNA-mRNA network, leading to activation of the Wnt and FoxO signaling pathways, ultimately promoting pulmonary hypertension^25,26^. Moreover, m^6^A-related single-nucleotide polymorphisms (m^6^A-SNPs) are crucial multifunctional genetic variants associated with cardiovascular disease and BP regulation^27,28^. The FTO-risk genotype, for example, exhibited higher sympathetic modulation of vasomotor tone and systolic BP^29^. Smoking increased BP by enhancing vascular FTO levels, thereby reducing m^6^A abundance and activating NOX2/ROS signaling pathway^30^. Additionally, FTO-induced erasure of m^6^A methylation has been reported to trigger vascular endothelial inflammation in diabetes^31^.

Endothelial dysfunction and hypertension are closely tied to NO bioavailability. The endothelial isoform of NO synthase (eNOS) is responsible for NO production in endothelium. eNOS is a well-known endothelium-derived relaxing factor that can be transcriptionally regulated or post-transcriptionally modified^32,33^. Our results align with m^6^A prediction of low confidence in *NOS3* from SRAMP, revealing that EC-METTL3 deficiency induces transcriptional downregulation without affecting its methylation level, suggesting that transcription factors may participate in the regulation of eNOS under METTL3-methylation. Factors such as laminar shear stress, TGF-beta1 and KLF2 have been reported to upregulate eNOS transcription^34^. While polymorphisms in the promoter region of eNOS are associated with coronary artery disease^35^. TNF-α suppresses the mRNA and protein levels of eNOS in systemic and pulmonary endothelium^36^. Further studies have shown that TNF-α can induce a decrease in eNOS promoter activity^37^. As a transcription factor, RUNX1 has been reported to closely linked with cardiovascular development and regulation of the function, and to propose as a potential therapeutic target for cardiovascular disease^12^, its relationship with eNOS has rarely been reported. RUNX1 score is high according to the Cistrome Data Browser prediction of NOS3 transcriptional regulation, indicating that NOS3 may be regulated by RUNX1. In current study, we identified RUNX1 as a novel eNOS promoter-binding transcription factor that downregulates eNOS mRNA levels. Therefore, RUNX1 negatively regulates the METTL3-eNOS cascade pathway. Runx1 knockout mice exhibit disrupted hematopoiesis and abnormal vasculature development, highlighting the important role of RUNX1 in development of vasculature^13^. RUNX1 deficiency protected against adverse cardiac remodeling after myocardial infarction. At present, RUNX1 is known to have four variants: RUNX1a, RUNX1b, RUNX1c, and RUNX1ΔN^19^. RUNX1a was not degraded by METTL3 but could be truncated at exon 7, as we and other teams have found^21^. RUNX1a can bind DNA but lacks the C-terminal transcriptional regulatory domain in RUNX1b, suggesting that RUNX1a may act as an antagonist to transcriptional activation compared with RUNX1b, thus explaining the different usage of RUNX1b and RUNX1a in RNA-seq and MeRIP-seq. Although RUXN1a is activated by METTL3, it is not involved in the transcriptional regulation of eNOS, while not excluding other vascular functions. In addition, the formation of RUNX1-CBFβ heterodimer protects RUNX1 from degradation by the ubiquitin ligase complex^38^. Ro5-3335 breaks the interaction between RUNX1 and CBFβ to inhibit the transcriptional activation^39^, then alleviates endothelial activation induced by enthelial-METLL3 deficiency.

In summary, our study provides compelling *in vitro* and *in vivo* evidence supporting the regulatory role of m^6^A RNA modification in the progression of endothelial activation in response to Ang II and subsequently elevates BP. Notably, METTL3-modified *RUNX1b* mRNA is involved in the pathogenesis of hypertension. The RUNX1b/eNOS axis emerges as a key contributor to m^6^A-modified hypertension, and inhibiting this axis holds promise for attenuating hypertension.

## Nonstandard Abbreviations and Acronyms

Ach: acetylcholine

ALKBH5: α-ketoglutarate-dependent dioxygenase alk B homolog 5

BP: blood pressure

ChIP: chromatin immunoprecipitation

ECs: endothelial cells

EDR: electron paramagnetic resonance

EPR: electron paramagnetic resonance

FTO: obesity-associated protein

L-NAME: L-NG-nitroarginine methyl ester

m^6^A: *N^6^*-methyladenosine

METTL3: methyltransferase-like 3

MeRIP–seq: m^6^A-specific methylated RNA immunoprecipitation combined with high-throughput sequencing

## Author contributions

B.L. and Y.Z. designed the research; B.L., Y.Z., M.L., Z.Y. and S.L. performed research; B.L., Y.Z., X.Y., Y.L., C.H., D.A. and Y.Y. analyzed data; B.L. and Y.Z. wrote the manuscript. All authors read and approved the final manuscript.

